# Excessive F-Actin and Microtubule Formation Mediates Primary Cilia Shortening and Loss in Response to Increased Extracellular Osmotic Pressure

**DOI:** 10.1101/2024.01.25.577175

**Authors:** Hiroshi Otani, Ryota Nakazato, Faryal Ijaz, Kanae Koike, Koji Ikegami

**Author notes:** The corresponding author: Koji Ikegami Department of Anatomy and Developmental Biology, Graduate School of Biomedical and Health Sciences, Hiroshima University, 1-2-3 Kasumi, Minami-Ku, Hiroshima 734-8553, Japan. Hiroshi Otani and Ryota Nakazato contributed equally to this work. **Abbreviations**: EV(s), extracellular vesicles; ICC, Immunocytochemistry; LatA, Latrunculin A; LIMK2, LIM domain kinase 2; PCM, pericentriolar material or pericentriolar matrix; PFA, paraformaldehyde; PI, propidium iodide; PLK1, polo-like kinase 1; PTX, paclitaxel; TAZ, transcriptional coactivator with PDZ-binding motif; TESK, testis associated actin remodeling kinase; TRP, transient receptor potential; YAP, Yes-associated protein.

## Abstract

The primary cilium is a small organelle protruding from the cell surface and is recognized as an antenna for signals from the extracellular milieu. Maintenance of primary cilia structure is crucial for proper behaviors of cells, tissues, and organs. While a dozen of studies have reported that several genetic factors impair the structure of primary cilia, evidence for environmental stimuli affecting primary cilia structures is limited. Here, we investigated an extracellular stress that affected primary cilia morphology and its underlying mechanisms. Hyperosmotic shock with increased extracellular sodium chloride concentration induced shortenings and disassembly of primary cilia in murine intramedullary collecting ducts cells. The shortening of primary cilia caused by hyperosmotic shock followed a loss of axonemal microtubules and delocalization of pericentriolar materials (PCMs). The primary cilia shortening/disassembly and PCMs delocalization were reversible. In parallel with these hyperosmotic shock-induced changes of primary cilia and PCMs, excessive microtubule and F-actin formation occurred in the cytoplasm. A microtubule-disrupting agent, Nocodazole, prevented the hyperosmotic shock-induced primary cilia disassembly partially, while preventing the delocalization of PCMs almost 100%. An actin polymerization inhibitor, Latrunculin A, also prevented partially the hyperosmotic shock-induced primary cilia shortening and disassembly, while preventing the delocalization of PCMs almost 100%. Taken together, we demonstrate that hyperosmotic shock induces reversible morphological changes in primary cilia and PCMs in a manner dependent on excessive formation of microtubule and F-actin.

## INTRODUCTION

Primary cilia, present in most vertebrate cell types, are tiny protruding organelles known to play critical roles as sensory antennae for extracellular signals (Singla and Reiter, 2006; Phua et al., 2015). Various receptors and channels concentrate on primary cilia and are responsible for mechanical and chemical receptions (Nauli et al., 2003; Rohatgi et al., 2007; Schou et al., 2015). Primary cilia function has been reported to be critical for cell proliferation and differentiation (Ezratty et al., 2011; Christensen et al., 2012). The structure of primary cilia is supported by well-organized bundles of microtubules, called axoneme. Axonemes of primary cilia are built on the mother centriole. The mother centriole has distal and subdistal appendages radially around its distal end, which create transition fibers to separate the ciliary space from cytoplasm (Sorokin, 1962; Loncarek and Bettencourt-Dias, 2018; Ma et al., 2023). The proximal ends of centrioles are surrounded by the pericentriolar matrix (PCM) (Woodruff et al., 2014), which consists of various proteins for organelle trafficking (Lee et al., 2021).

The morphology of primary cilia depends on the balance among ciliogenesis, ciliary elongation, and ciliary resorption. Ciliogenesis starts with the assembly of a ciliary vesicle at the distal end of the mother centriole (Lu et al., 2015; Blanco-Ameijeiras et al., 2022). Both distal and subdistal appendages are required for the cilia-directing vesicular transport and ciliary vesicle formation to initiate ciliogenesis (Čajánek and Nigg, 2014). Tubulin dimers subsequently polymerize to form axonemal microtubules at the distal end of mother centriole. This process requires pericentriolar proteins that constitute PCMs including ODF2 (Tateishi et al., 2013). The centrioles are docked to the cell membrane and the polymerizing axoneme protrudes from the cell surface (Garcia-Gonzalo and Reiter, 2012; Reiter et al., 2012). Ciliary elongation depends on anterograde intraflagellar transport (IFT) of ciliary materials. The ejection of ciliary components from primary cilia by retrograde IFT and destabilization of axonemes by deacetylation of tubulin in primary cilia causes resorption of primary cilia (Breslow et al., 2013; Lin et al., 2013; Phua et al., 2017).

The balance among ciliogenesis, ciliary elongation, and ciliary resorption is changed by cellular states. Ciliogenesis occurs during the non-dividing phase of the cell cycle, while ciliary resorption occurs under the mitotic phase (Plotnikova et al., 2009). Cell-intrinsic circadian rhythms also regulate ciliary elongation and resorption (Tu et al., 2023; Nakazato et al., 2023a). Resorption of primary cilia occurs in response to some stimulation as well. Serum stimulation induces extracellular vesicle release from primary ciliary tips, which triggers rapid loss of IFT-B from primary cilia and resorption of primary cilia (Phua et al., 2017). Heat shock from elevated extracellular temperatures induces resorption of primary cilia mediated by a deacetylase HDAC6 (Prodromou et al., 2012).

Cytoskeletal microtubule and actin filament dynamics affect the morphology of primary cilia. The dynamics of microtubules in the cytoplasm as well as axonemes in primary cilia are also known to contribute significantly to the formation of primary cilia. It has been reported that the amount of free tubulin dimmers in the cytoplasm regulates the length of primary cilia; microtubule stabilization shortens the length of primary cilia, while accelerated microtubule depolymerization elongates them (Sharma et al., 2011). Docking of the mother centrioles with ciliary vesicles to the cell membrane, which is observed in the early stages of ciliogenesis, requires the removal of cortical actin filaments (Jewett et al., 2021; Tanos et al., 2013). Actin remodeling factors, including LIMK2 and TESK1, were reported to control ciliogenesis by regulating the transcriptional coactivator Yes-associated protein (YAP)/transcriptional coactivator with PDZ-binding motif (TAZ) and directional ciliary vesicle trafficking (Kim et al., 2015). Migration of centrosome to the apical surface during ciliogenesis in polarized epithelial cells depends on the microtubule network densification and the actin cytoskeleton asymmetrical contraction (Pitaval et al., 2017). Actin polymerization within primary cilia also contributes to ciliary resorption, following scission of ciliary tips (Phua et al., 2017).

Unlike normal body fluid osmolarity (≈300 mOsm/kg), urine osmolarity fluctuates ranging from 50 to 1,200 mOsm/kg due to renal reabsorption (Koeppen and Stanton, 2012; Kitiwan et al., 2021). Primary cilia are also present on the epithelial cells of renal collecting ducts. Primary cilia protruding toward the lumen of the collecting ducts are therefore exposed to a wide range of osmolarity changes caused by urine (Webber and Lee, 1975). Nonetheless, the effects of osmotic alteration on primary cilia are poorly understood. Here, we investigated the impact of hyperosmotic shock on the morphology of primary cilia, their supporting structures in epithelial cells of renal collecting ducts, and their underlying mechanisms using cytoskeleton-targeting reagents.

## RESULTS

### Acute hyperosmotic shock shortens and disassembles the primary cilia of mIMCD-3 cells

To determine whether extracellular stress impacts the shape of primary cilia, we investigated the effect of hyperosmotic shock on a cell line derived from murine intramedullary collecting ducts epithelial cells, mIMCD-3 cells. The medium osmolarity we used was labeled ≈310 mOsm/kg (isoosmotic; NaCl: 120 mM; NaHCO_3_: 29 mM; Osmolarity: 295–325 mOsm/kg). By adding different amounts of 5M NaCl to this isoosmotic medium, we exposed mIMCD-3 cells to hyperosmotic shock ranging from ≈410 to 810 mOsm/kg for 3 h. We visualized primary cilia by staining ARL13B, a primary ciliary marker, and the ciliary base by staining γ-tubulin, a pericentriolar protein, with immunocytochemistry (ICC) (Fig. 1A).

**Figure 1.**
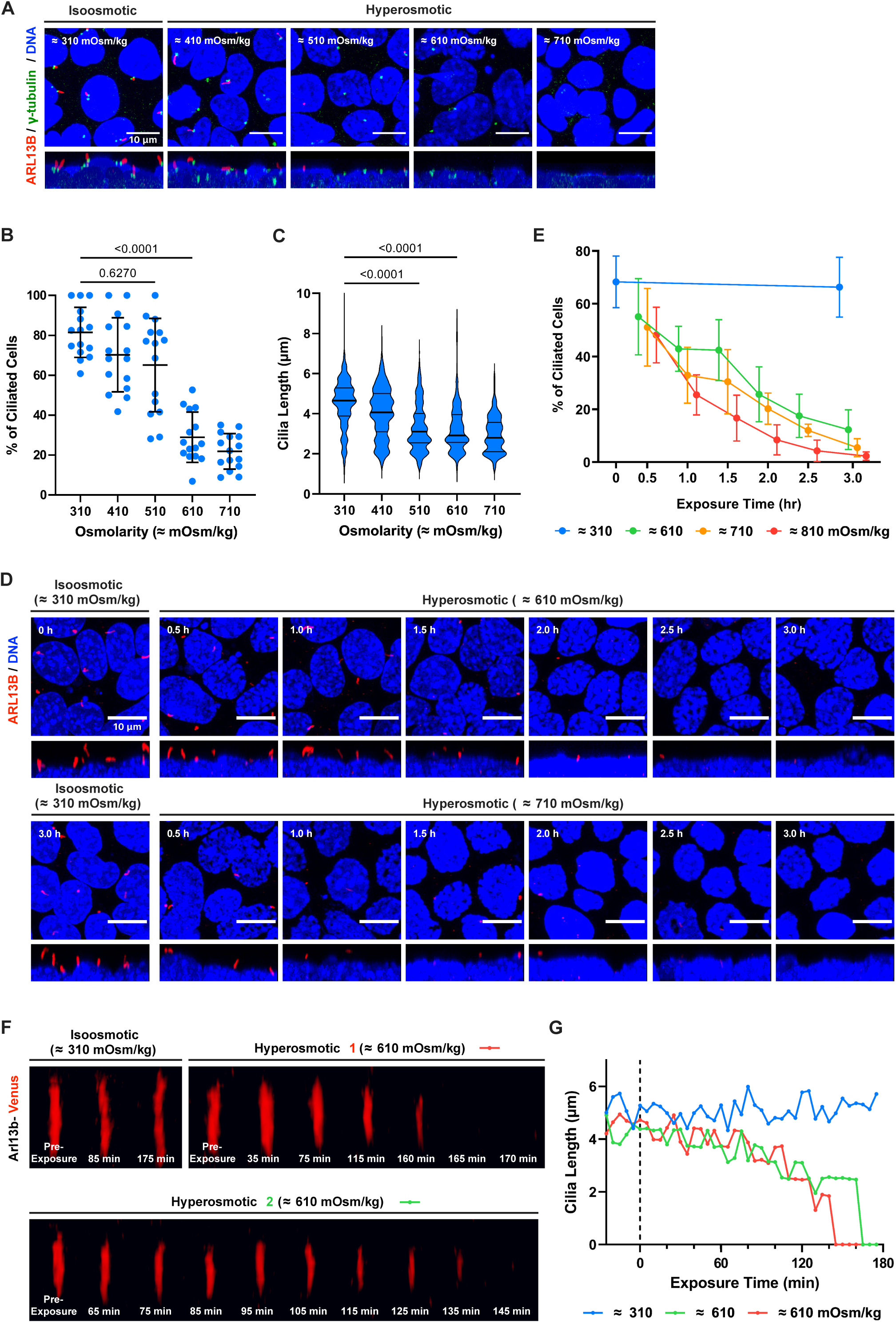
Acute hyperosmotic shock shortens and disassembles the primary cilia of mIMCD-3 cells in a time-dependent manner. **(A)** Confocal images of mIMCD-3 after immunocytochemistry (ICC) for primary cilia (red, ARL13b), pericentriolar protein as the base of primary cilia (green, γ-tubulin), and DNA (blue, DAPI). Ciliated cells were incubated at the indicated osmolarity for 3 h. Scale bars: 10 µm. (B) Quantification of the percentage of ARL13B-positive cilia-possessing cells in A. The data are from five fields per condition from three independent experiments. Bars show mean ± SD. (C) Quantification of ciliary lengths on the cells in A. The data are from 15 fields per condition from three independent experiments. Bars show median ± quartile deviation. (D) Confocal images of time-coursed ICC samples. Ciliated cells were incubated at the indicated osmolarity for the indicated time periods. Scale bars: 10 μm. (E) Quantification of the percentage of ARL13B-positive cells in D. The data are from five fields per condition from three independent experiments. Error bars show mean ± SD. (F) Rendered three-dimensional images of representative primary cilia acquired from the confocal live imaging on ciliated Arl13b-Venus-expressing mIMCD-3 cells at indicated osmolarity. Time stamps indicate the time that passed since hyperosmotic shock started. (G) Quantification of the length of primary cilia in F.

Under the normal medium osmolarity, the ARL13B-positive primary cilia were observed in 81.5 ± 13.0% of total cells (Fig. 1B). Under the hyperosmotic shock, the number of ARL13B-positive primary cilia was decreased compared with the isoosmotic conditions, dependently on the increase of the extracellular osmolarity (Fig. 1A and 1B). Cells under hyperosmotic shock at higher than ≈610 mOsm/kg exhibited statistically significant loss of primary cilia; 28.9 ± 12.6% of cells possessed ARL13B-positive primary cilia in ≈610 mOsm/kg culture medium and only 21.9 ± 8.9% in ≈710 mOsm/kg medium (Fig. 1A and 1B). Under milder hyperosmotic shock at ≈410 or ≈510 mOsm/kg, the primary cilia-positive rate decreased less (Fig. 1B; ciliated cells: 70.2 ± 18.5 % in ≈410 mOsm/kg, 65.1 ± 23.4 % in ≈510 mOsm/kg).

We quantified the effects of hyperosmotic shock on the length of the remaining primary cilia. The mean length of primary cilia in control cells was 4.5 ± 1.2 µm (Fig. 1C). Under the hyperosmotic shock, the mean length of primary cilia became shorter (Fig. 1A and 1C). In contrast to the ARL13B-positive primary cilia rate, the length of remaining primary cilia in ≈510 mOsm/kg was significantly shorter than in the isoosmolarity (Fig. 1C). Furthermore, in higher osmolarity, the length of the remaining primary cilia was further reduced (Fig. 1C). These data indicate that in epithelial cells of collecting ducts, milder hyperosmotic shock shortens the length of primary cilia, and sever hyperosmotic shock causes the loss of primary cilia.

To track the disassembly of the primary cilia in a time course, we performed the ICC of primary cilia by staining ARL13B in mIMCD-3 cells at different exposure times of hyperosmotic shock; 0.5, 1.0, 1.5, 2.0, 2.5, and 3.0 h (Fig. 1D). The number of ARL13B-positive primary cilia in cells exposed to hyperosmotic shock decreased gradually in a time-dependent manner, with a statistically significant decrease after 1 h in ≈810 mOsm/kg culture medium (25.5 ± 7.6%) and after 2 h in ≈710 mOsm/kg (20.1 ± 5.9%) and ≈610 mOsm/kg culture medium (25.7 ± 10.4%) (Fig. 1E).

We next carried out live cell imaging to observe how primary cilia shortened and disassembled during the 3 hours of hyperosmotic shock. Since overexpressing ciliary marker proteins causes abnormal shapes and elongation in primary cilia (Nachury et al., 2007; Ijaz and Ikegami, 2021), we chose a cell line that expresses stably and endogenously Arl13b-Venus. To eliminate unknown effects of knock-in antibiotics-resistant gene expression, we directly inserted Venus into the C-terminus of endogenous Arl13b with the HITI method. With the HITI method, we generated a clone where an allele of Arl13b was C-terminally-tagged with Venus. We performed the three-dimensional time-lapse imaging of the Arl13b-Venus-expressing mIMCD-3 cells under isoosmolarity (≈310 mOsm/kg) and hyperosmolarity (≈610 mOsm/kg) for 3 h respectively (Movie S1, Fig. 1F). In the acquired movie, almost half of the primary cilia got shorter over time and most of them eventually disappeared after the 3 h of hyperosmotic shock, ≈610 mOsm/kg (Movie S1, right; Fig. 1F, Hyperosmotic). In contrast, the majority of primary cilia retained their length throughout the 3 h of recording time under the isoosmolarity, ≈310 mOsm/kg (Movie S1, left; Fig. 1F, Isoosmotic), indicating that the laser emission from the confocal microscope had little effect on the length of primary cilia or the fluorescence intensity of Venus. We also measured the length of primary cilia at each time point in the time-lapse data (Fig. 1G). Primary cilia kept their length in the first hour and began to shorten after one hour of exposure to the hyperosmotic shock (Fig. 1G; 0-60 min). The shortenings accelerated in the second hour (Movie S1, Fig. 1G; 60-120 min), and the shortened primary cilia finally vanished in several minutes during the third hour (Movie S1, Fig. 1G; 120-180 min). This time-lapse imaging demonstrates that hyperosmotic shock induces the gradual shrinking and consequent loss of primary cilia.

### Acute hyperosmotic shock shortens and disassembles the axoneme prior to ciliary membrane shortening

Figure 1 shows the shortening and loss of primary cilia with increase in the amount of hyperosmotic shock. ARL13B was used as a marker for primary cilia. It was unclear whether the ciliary axoneme behaved the same way as the ciliary membrane. To examine whether the ciliary axoneme is affected by hyperosmotic shock, we exposed cells to hyperosmotic shock at ≈510 or ≈710 mOsm/kg for 3 h and performed ICC on both the ciliary axoneme (α-tubulin) and ARL13B (Fig. 2A). We then quantified the lengths of remaining primary cilia positive for α-tubulin and/or for ARL13B, and plotted them on 2D diagrams (Fig. 2B). The linear regression line drawn between the α-tubulin-positive cilia length and the ARL13B-positive cilia length represents the correlation between the ciliary membrane and the ciliary axoneme. The shallower slope of the linear regression line represents the shorter ciliary axoneme length (α-tubulin-positive cilla length) than the ARL13B length (ARL13B-positive cilia length). We also quantified the percentage of α-tubulin-positive cilia (Fig. 2C). With hyperosmotic shock at ≈510 mOsm/kg, the slope of the linear regression line was shallower (y = 0.3034x), and the percentage of α-tubulin-positive cilia was lower (27.6 ± 18.6%) than those with isoosmolarity (y = 0.616x; 81.2 ± 11.2%). Interestingly, the slope of the linear regression line and the percentage of α-tubulin-positive cilia with hyperosmotic shock at ≈710 mOsm/kg (y = 0.5132x; 57.0 ± 24.1%) were higher than those with hyperosmotic shock at ≈510 mOsm/kg (Fig. 2B and 2C). These data indicate that the ciliary axoneme gets resorbed, preceding ciliary membrane shortening, with hyperosmotic shock.

**Figure 2.**
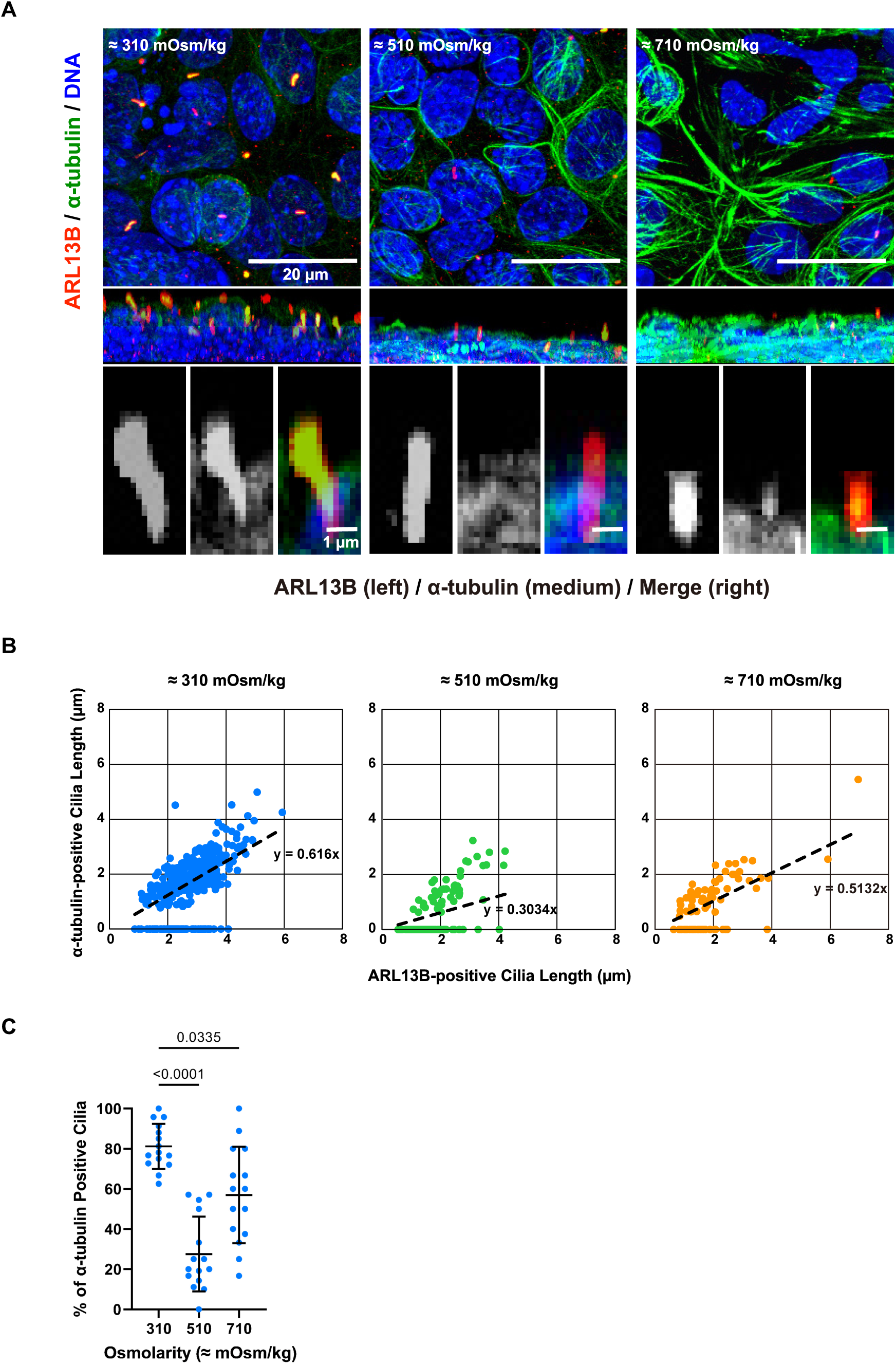
Acute hyperosmotic shock shortens and disassembles the axoneme. **(A)** Confocal images of mIMCD-3 cells after ICC for primary cilia (red, ARL13B), microtubules (green, α-tubulin), and DNA (blue, DAPI). Cells were incubated at the indicated osmolarity for 3 h. Scale bars: 20 μm. (B) Quantification and 2-D plot of α-tubulin-positive cilia length and ARL13B-positive cilia length in A. The dashed line with equations represents the linear regression of the plots. (C) Quantification of the percentage of α-tubulin-positive cilia against ARL13-positive cilia in A. The data are from five fields per condition from three independent experiments. Error bars show mean ± SD.

### Hyperosmotic shock disrupts the pericentriolar matrix

We next investigated whether pericentriolar materials (PCMs) disrupted under hyperosmotic shock. We immunostained two pericentriolar proteins, ODF2 and γ-tubulin, as well as DNA after 3 h of hyperosmotic shock at ≈510, ≈610, ≈710, and ≈810 mOsm/kg (Fig. 3A). ODF2 and γ-tubulin showed different responses against hyperosmotic shocks. Although the positive rates of both proteins decreased in a hyperosmotic shock-dependent manner, γ-tubulin showed more tolerance against hyperosmotic shock than ODF2 (Fig. 3B and 3C). While the ODF2-positive rate significantly dropped at ≈610 mOsm/kg with the lowest rate at ≈710 mOsm/kg (Fig. 3B), the γ-tubulin-positive rate showed a statistically significant drop at ≈710 mOsm/kg or more (Fig. 3C).

**Figure 3.**
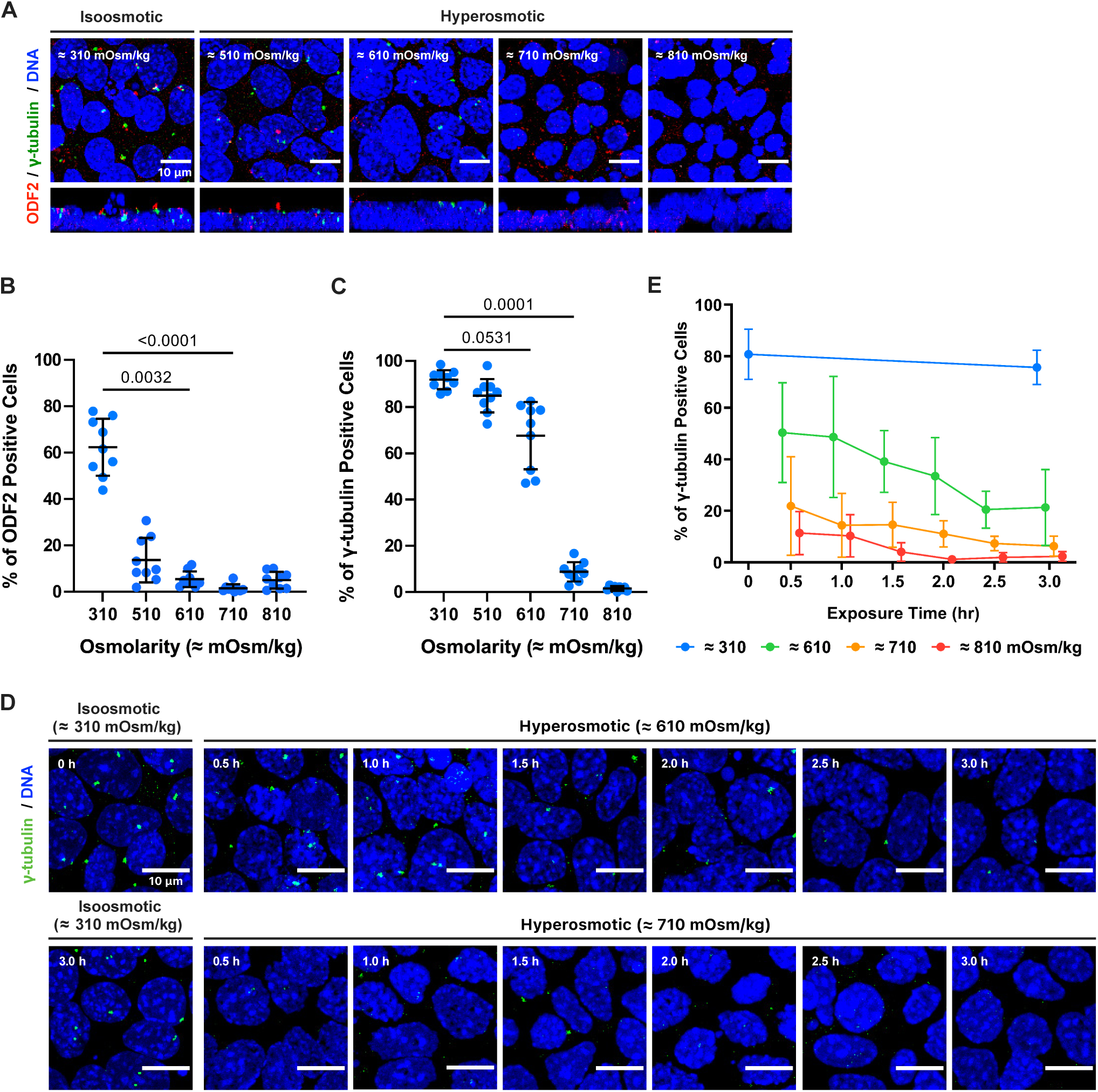
Hyperosmotic shock disrupts the pericentriolar matrix (PCM). **(A)** Confocal images of mIMCD-3 cells after ICC for two pericentriolar proteins (red, ODF2; green, γ-tubulin) and DNA (blue, DAPI). Ciliated cells were incubated at the indicated osmolarity for 3 h. Scale bars: 10 µm. (**B and C**) Quantifications of the percentage of ODF2-positive (B) and γ-tubulin-positive cells (C) in A. The data are from five fields per condition from three independent experiments. Bars show mean ± SD. **(D)** Confocal images of time-coursed ICC samples. Ciliated cells were incubated at the indicated osmolarity for the indicated time periods. Scale bars: 10 μm. (E) Quantification of the percentage of γ-tubulin-positive cells in D. The data are from five fields per condition from three independent experiments. Error bars show mean ± SD.

We further observed how the γ-tubulin-positive rate decreased in a time course at different hyperosmolarity as we did on primary cilia. We immunostained γ-tubulin in mIMCD-3 cells after different exposure times of hyperosmotic shock (Fig. 3D). Out of all the hyperosmolarity we tested, the γ-tubulin-positive rate dropped throughout the 3 h of hyperosmotic shock (Fig. 3D). At ≈710 and ≈810 mOsm/kg, the highest decrease occurred between 0 to 0.5 h time points at both hyperosmolarity (Fig. 3E; 21.8 ± 19.1% in ≈710 mOsm/kg medium and 11.3 ± 8.4% in ≈810 mOsm/kg medium). These were highly contrasting to the results of ARL13B-positive cilia resorption; about 50% ARL13B-positive primary cilia were still present at 0.5 h after exposure to ≈710 or ≈810 mOsm/kg hyperosmotic shock (Fig. 1E). These results demonstrate that pericentriolar proteins, including γ-tubulin and ODF2, disappear from the base of primary cilia prior to the primary cilia shortening and disassembly during hyperosmotic shocks.

### Loss of primary cilia and PCM markers resulted from neither shock due to acute change of osmolarity, protein degradation, nor disassembly of centriole core

Osmolarity difference in cell membrane creates sheer stress on cells, especially on the structures with high surface area-to-volume ratio like primary cilia (William, 2005). We aimed to exclude the possibility of primary cilia to be torn apart by the physical and osmotic stress we have tested. We incubated the cells at three different temperatures: 4, 22, and 37 °C during the 3-h hyperosmotic shock at ≈610 and ≈710 mOsm/kg. We immunostained ARL13B to observe primary cilia (Fig. S1A). At isoosmolarity, the cilia-positive rate did not significantly change by temperature (Fig. S1B). After hyperosmotic shock at ≈610 and ≈710 mOsm/kg, the cilia-positive rate decreased significantly at 37 °C (Fig. S1C, S1D). With hyperosmotic shock at ≈610 mOsm/kg, the cilia-positive rate dropped as low as 10.7 ± 0.9% at 37 °C compared to the 65.5 ± 4.6% at 4 °C and 66.1 ± 2.2% at 22 °C (Fig. S1C). These results suggest that the hyperosmotic shock-induced primary cilia loss is not caused by sheer physical and osmotic stress, but by the enzymatic and biochemical processes.

The method of hyperosmotic exposure used gave a steep rate of osmolarity increase especially when the osmolarity was ≈610 or ≈710 mOsm/kg (Fig. 1, 2). To exclude a possibility of the cilia shortening/disassembly and PCMs disruption being artifacts caused by the acute change of osmolarity, we exposed cells to gradual increase in medium osmolarity; ≈410 mOsm/kg for 1 h, ≈510 for 1 h, and ≈610 for 2 h (Fig. 4A). The gradual increase of osmolarity induced the same morphological changes (Fig. 4B); ARL13B-positive primary cilia loss (Fig. 4C; from 75.4 ± 11.2 to 7.7 ± 4.8%) and the γ-tubulin loss (Fig. 4D; from 92.4 ± 4.9 to 57.4 ± 16.5%). These indicate that the hyperosmotic shock-dependent loss of ARL13B-positive primary cilia and PCMs does not result from acute increase in the medium osmolarity.

**Figure 4.**
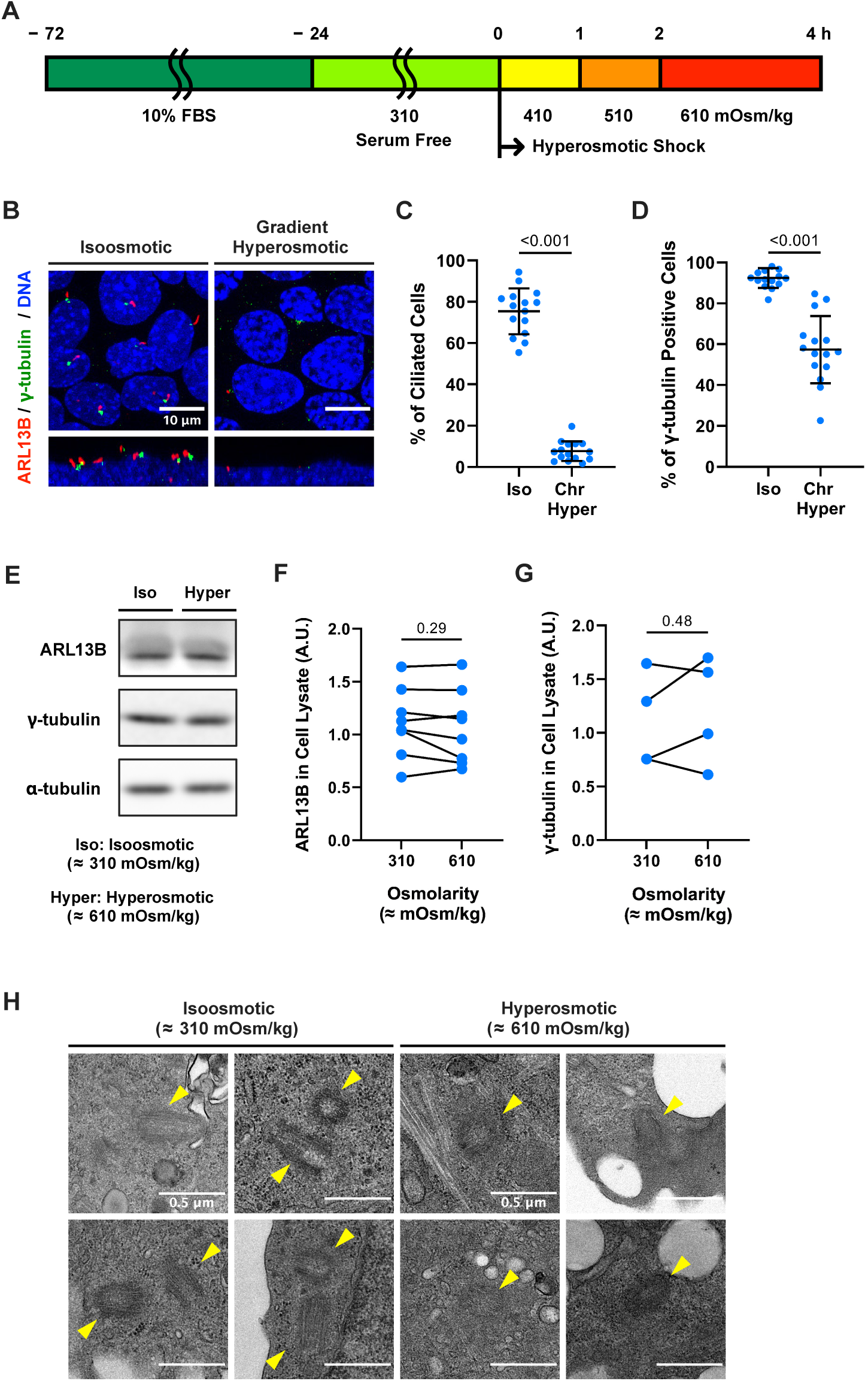
Loss of primary cilia and PCM markers resulted from neither shocks due to acute change of osmolarity, protein degradation, nor disassembly of centriole core. **(A)** Schematic diagram for the gradual increase of medium osmolarity. Cells were exposed to medium osmolarity ≈410 mOsm/kg for 1 h, ≈510 mOsm/kg for 1 h, and ≈610 mOsm/kg for 2 h. (B) Confocal images of mIMCD-3 after ICC for primary cilia (red, ARL13B), γ-tubulin (green), and DNA (blue, DAPI). Scale bars: 10 µm. (C and D) Quantification of the percentage of ARL13B-positive cilia-possessing cells (C) and the percentage of γ-tubulin-positive cells (D). Bars show mean ± SD. (E) Western blot analyses of expression of ARL13B and γ-tubulin in mIMCD-3 cells at each indicated osmolarity at 3 hours after hyperosmotic shock started. (F and G) Quantification of the expression level of ARL13B (F) and γ-tubulin (G) in E. (H) Transmission electron microscopy (TEM) of mIMCD-3 cells fixed after 3 h exposure to the indicated osmolarity. Yellow arrows point to centrioles. Scale bars: 0.5 μm.

To examine whether degradation of cilia- and PCMs-related proteins caused the cilia shorteing/loss and PCMs disruption, we investigated the amount of ARL13B and γ-tubulin in the cell lysate with western blot analyses after the 3 h of hyperosmotic shock at ≈610 mOsm/kg (Fig. 4E). ARL13B, a cilia-related protein, showed almost no significant change in its protein level (Fig. 4F), suggesting that the hyperosmotic shock-induced cilia loss neither necessarily affect the cilia-related protein level nor result from ciliary proteins degradation. While both γ-tubulin and ODF2 disappeared by the hyperosmotic shock in ICC, γ-tubulin did not show a statistically significant change in protein level (Fig. 4G), suggesting that hyperosmotic shock does not degrade the pericentriolar proteins but could delocalize them.

As the hyperosmotic shock leads to PCM disruption in ICC, we performed transmission electron microscopy (TEM) on centrioles to examine whether the centrioles are not disassembled by hyperosmotic shock at ≈610 mOsm/kg (Fig. 4H). The representative images show that the centrioles are not disassembled with hyperosmotic shock. In contrast, the electron density around the centrioles in hyperosmotic shock is less than that of isoosmolarity (Fig. 4H). These results further indicate hyperosmotic shock-induced delocalization of PCMs.

### Hyperosmotic shock-induced cilia loss and PCM delocalization are reversible

To clarify whether hyperosmotic shock-induced cilia loss and PCM disruption do not result from cell death, we stained the cells with DAPI and propidium iodide (PI) immediately after 3 h of hyperosmotic shock (Fig. 5A). Even under the highest osmolarity we tested, ≈810 mOsm/kg, the cell death rate was below 10% (Fig. 5B). The osmolarity that was enough to induce cilia loss and PCM disruption, ≈610 mOsm/kg, leaded to almost no cell death (Fig. 5B).

**Figure 5.**
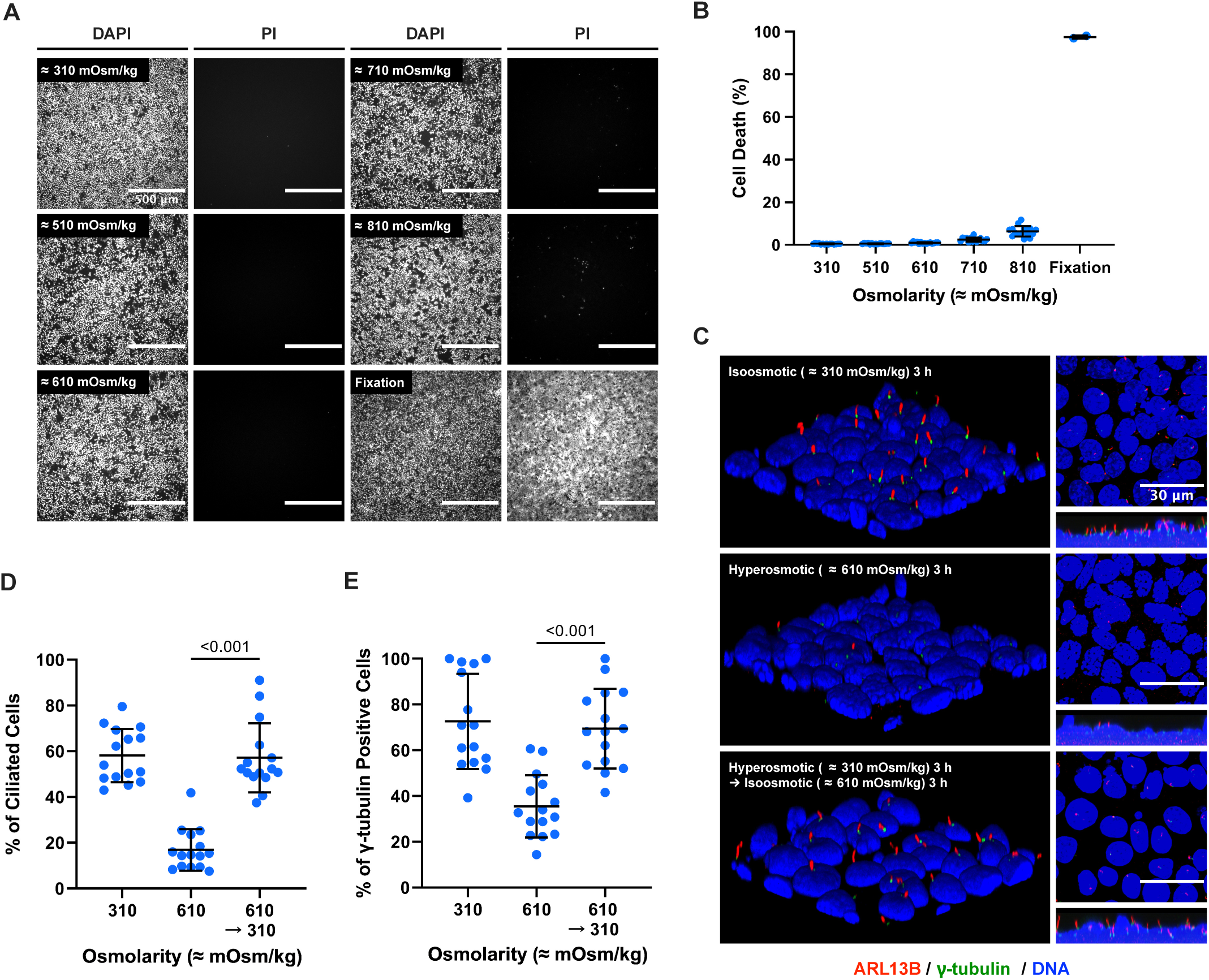
Hyperosmotic shock-induced cilia loss and PCM delocalization are reversible. **(A)** Fluorescence microscopy of mIMCD-3 cells stained for dead cells (red, propidium iodide: PI) and all cells (blue, DAPI) in indicated osmolarity at 3 hours after hyperosmotic shock started. Cells fixed with 4% PFA were shown as a positive control (Fixation). Scale bars: 500 µm. (B) Quantification of the percentage of PI-positive cells in A. The data are from five fields per condition from three independent experiments. Bars show mean ± SD. (C) Confocal images of mIMCD-3 cells after ICC for primary cilia (red, ARL13B), pericentriolar protein (green, γ-tubulin), and DNA (blue, DAPI). Ciliated mIMCD-3 cells were incubated in isosmotic condition (upper panel) or hyperosmotic condition (middle panel) for 3 h, or hyperosmotic condition for 3 h followed by incubation in isosmotic condition for additional 3 h (bottom panel). Scale bars: 30 µm. (D and E) Quantification of the percentage of ARL13B-positive cilia-possessing cells (D) and γ-tubulin-positive cells (E) in C. The data are from five fields per condition from three independent experiments. Bars show mean ± SD.

To further investigate the lasting effect of the hyperosmotic shock and examine whether the morphological changes of the primary cilia and the PCM are reversible, we restored the osmolarity of the medium to isoosmotic, ≈310 mOsm/kg, and incubated the cells for another 3 h after the 3-h hyperosmotic shock at ≈610 mOsm/kg, followed by staining ARL13B and γ-tubulin with ICC (Fig. 5C). Both the primary cilia-positive rate and the γ-tubulin-positive rate were restored close to that before the hyperosmotic exposure; the primary cilia-positive rate was 58.1% before the hyperosmotic shock and increased back to 57.1% at 3 h after the medium osmolarity was restored to isoosmotic (Fig 5D). The γ-tubulin-positive rate was 72.6% before the hyperosmotic shock and 69.4% after the osmolarity restoration (Fig 5E). Because the hyperosmotic shock we tested did not cause significant cell death, the primary cilia disassembly and the loss of the PCM are temporal phenomena and are reversible without taking longer than the cell cycle.

### Inhibition of microtubule polymerization reduces the hyperosmotic shock-induced primary cilia shortening/disassembly and the PCM delocalization

Next, we applied pharmacological approaches to investigate how cells shorten or disassemble their primary cilia. Our findings showed hyperosmotic shock caused loss of axonemal microtubules preceding ARL13B-positive ciliary membrane loss while inducing excessive formation of cytoplasmic microtubule bundles (Fig. 2). We thus investigated, as a mechanistic insight, contribution of microtubule to the shortening/loss of primary cilia and delocalization of PCMs by hyperosmotic shock. We first tested the effects of Nocodazole, which promotes microtubule depolymerization, on the primary cilia shortening/loss and PCMs delocalization. We exposed cells to hyperosmotic shock (≈610 mOsm/kg) for 3 h with or without Nocodazole. Nocodazole at 400 nM strongly blocked the hyperosmotic shock-induced excessive formation of cytoplasmic microtubule bundles (Fig. 6A). Under this condition, we performed ICC of ARL13B, γ-tubulin, and ODF2 (Fig. 6B). Remarkably, the microtubule polymerization inhibitor prevented the hyperosmotic shock-induced primary cilia loss (Fig. 6B; upper). The percentage of ciliated cells with isoosmolarity was ≈70% and decreased to ≈20% with hyperosmolarity (≈610 mOsm/kg) without Nocodazole; the rate got recovered to ≈50 % with 400 nM of Nocodazole (Fig. 6C). Nocodazole prevented delocalization of PCMs more effectively (Fig. 6B; middle and bottom). The percentage of γ-tubulin-positive cells and ODF2-positive cells got recovered to the same level of isoomolarity control (Fig. 6E and 6F). These results indicate that the hyperosmotic shock-induced morphological changes of primary cilia and delocalization of PCMs are dependent on excess microtubule formation.

**Figure 6.**
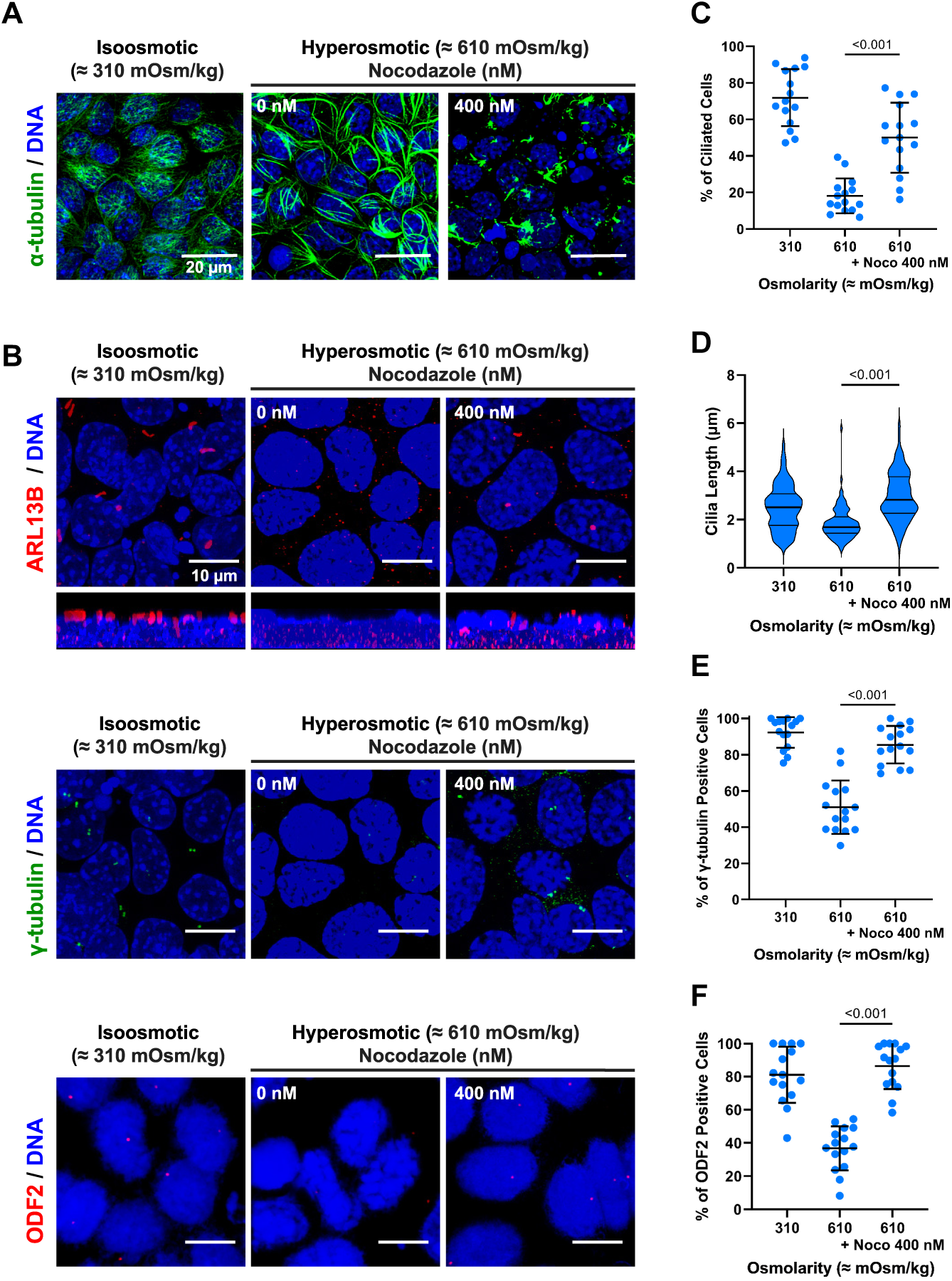
Inhibition of microtubule polymerization reduces the hyperosmotic shock-induced primary cilia shortening/disassembly and the PCM delocalization. **(A)** Confocal images of mIMCD-3 cells after staining for microtubules (green, α-tubulin) and DNA (blue, DAPI). Cells were exposed to the indicated osmolarity and Nocozole with DMSO as a vehicle. Scale bars: 20 μm. **(B)** Confocal images of mIMCD-3 cells after ICC for primary cilia (red, ARL13B, upper) or pericentriolar proteins (green, γ-tubulin, middle; red, ODF2, bottom), and DNA (blue, DAPI). Cells were exposed to the indicated osmolarity and Nocodazole with DMSO as a vehicle. Scale bars: 10 μm. (C) Quantification of the percentage of the ciliated cells in B. Bars show mean ± SD. (D) Quantification of the primary ciliary length of the ciliated cells in B. Bars show median ± quartile deviation. (E and F) Quantification of the percentage of the γ-tubulin-positive cells (E) and ODF2-positive cells (F) in B. Bars show mean ± SD.

Examination with paclitaxel (PTX), a microtubule polymerization-promoting agent, further supported the idea that the hyperosmotic shock-induced primary cilia shortening/loss occurred upon excessive microtubule formation. Excess formation of microtubule bundles in cytoplasm by PTX was confirmed by staining microtubules with an anti-α-tubulin antibody (Fig. S2A). PTX failed to prevent the hyperosmotic shock-induced loss of primary cilia at any PTX concentration we tested (Fig. S2B and S2C). The lengths of remaining primary cilia in the hyperosmolarity were also not recovered by any concentrations of PTX tested (Fig.S2D).

### Inhibition of F-actin formation reduces the hyperosmotic shock-induced primary cilia shortening/disassembly and the PCM delocalization

We next explored contribution of F-actin to the shortening/loss of primary cilia and delocalization of PCMs by hyperosmotic shock. To this end, we tried Latrunculin A (LatA), an inhibitor for F-actin formation. The effect of LatA on F-actin formation was confirmed by labeling cells with fluorescent phalloidin. LatA effectively disrupted F-actin in a concentration-dependent manner under normal condition (Fig. 7A), and suppressed excess formation of F-actin by hyperosmotic shock (Fig. 7B). The hyperosmotic shock-induced primary cilia loss was partially prevented by LatA even at 200 nM (Fig. 7C; red). Rate was kept as high as ≈40% with 200 nM LatA while the cilia-positive rate decreases from ≈80% to ≈21% by hyperosmotic shock without LatA (Fig. 7C and 7D). Even with the highest concentration of LatA, 1000 nM, the loss of primary cilia by hyperosmotic shock was not completely prevented; 1000 nM LatA held the cilia-positive rate at ≈50% under the hyperosmotic shock (Fig. 7D). Also, LatA partially prevented the hyperosmotic shock-induced shortening of the length of primary cilia at any LatA concentration; the hyperosmotic shock-induced primary cilia shortening was not completely prevented even with the highest concentration of LatA, 1000 nM (Fig. 7E).

**Figure 7.**
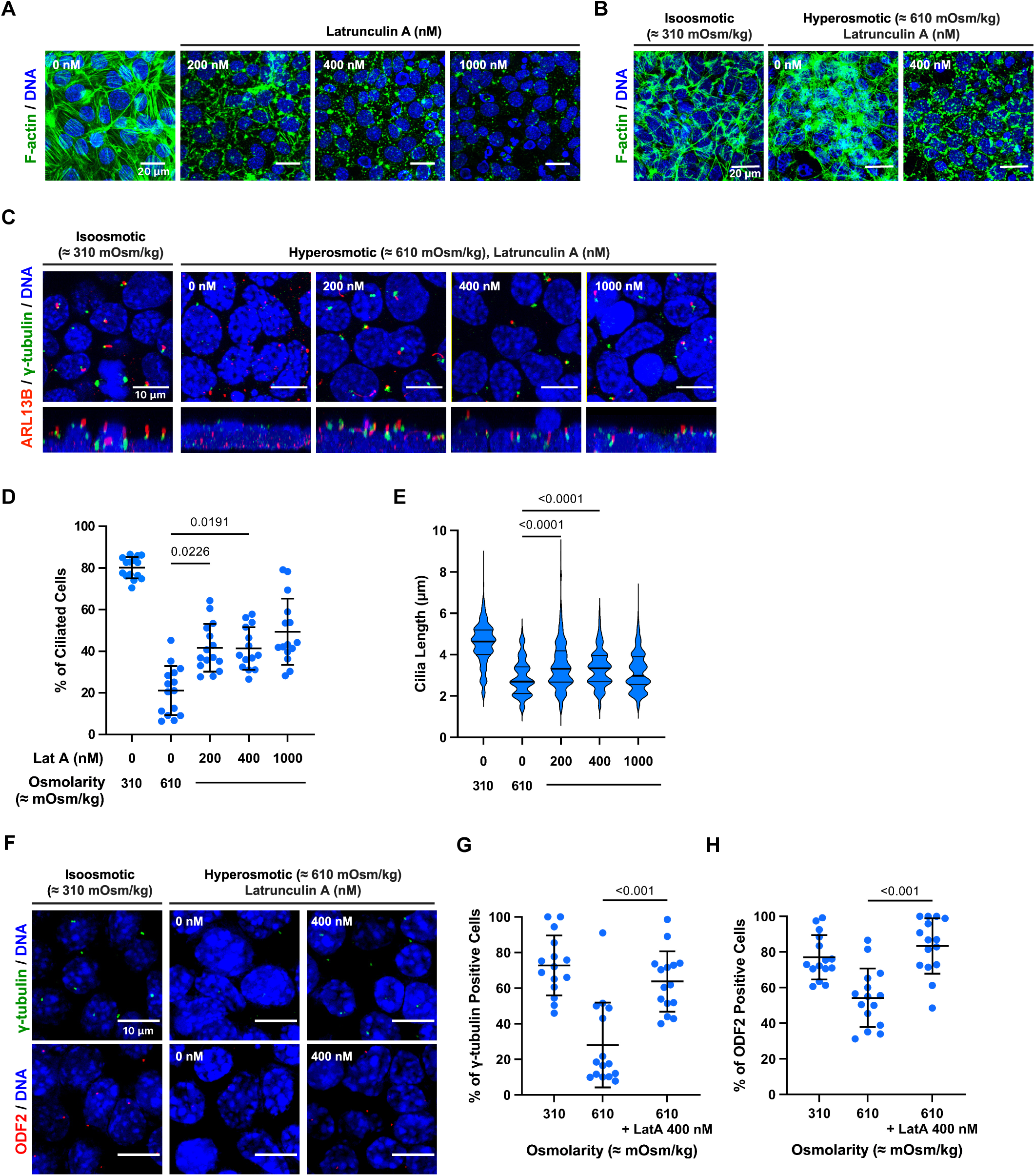
Inhibition of F-actin formation reduces the hyperosmotic shock-induced primary cilia shortening/disassembly and the PCM delocalization. **(A)** Confocal images of mIMCD-3 cells after staining for F-actin (green). Cells exposed to the indicated concentration of Latrunculin A (LatA) or DMSO as a vehicle. Scale bars: 20 µm. **(B)** Confocal images of mIMCD-3 cells after staining for F-actin (green). Cells were exposed to the indicated osmolarity and Latrunculin A or DMSO as a vehicle. Scale bars: 20 µm. **(C)** Confocal images of mIMCD-3 cells after ICC for primary cilia. The indicated concentration of LatA was added to hyperosmotic media, and DNA (blue), ARL13B (red), and γ-tubulin (green) were stained with ICC. Scale bars: 10 µm. (D) Quantification of the percentage of ARL13B-positive cilia-possessing cells in C. The data are from five fields per condition from three independent experiments. Bars show mean ± SD. (E) Quantification of ciliary lengths on the cells in C. The data are from 15 fields per condition from three independent experiments. Bars show median ± quartile deviation. **(F)** Confocal images of mIMCD-3 cells after ICC for two pericentriolar proteins. The indicated concentration of LatA was added to hyperosmotic media, and DNA (blue), ODF2 (red), and γ-tubulin (green) were stained with ICC. Scale bars: 10 µm. (G and H) Quantifications of the percentage of γ-tubulin-positive cells (G) and ODF2-positive cells (H) in F. The data are from five fields per condition from three independent experiments. Bars show mean ± SD.

To investigate whether LatA also prevents the delocalization of the PCMs, we stained the pericentriolar proteins, γ-tubulin and ODF2, after 3 h of hyperosmotic shock (≈610 mOsm/kg) with 400 nM LatA present (Fig. 7F). The rates of γ-tubulin- or ODF2-positive cells were recovered significantly by LatA treatment under hyperosmotic shock (Fig. 7G and 7H). Combined with the prevention of hyperosmotic shock-induced cilia shortening/loss by LatA, these data suggest that both the hyperosmotic shock-induced morphological changes of cilia shortening/loss and PCMs delocalization are F-actin-dependent.

### Hyperosmotic shock increases the amount of cilia fragments in the culture medium

We finally examined whether hyperosmotic shock-induced primary cilia shortening/loss associates with excision of primary cilia tips, given that the hyperosmotic shock-induced primary cilia shortening/loss was F-actin-dependent. We collected extracellular vesicle-like particles from the conditioned media by ultracentrifugation after 3 h of hyperosmotic shock to measure the amount of primary cilia fragments released into the cultured media. ARL13B and IFT88 were detected in the conditioned media pellet as well as in the cell lysate by western blot analyses (Fig. 8A). Interestingly, the two ciliary proteins showed opposite release tendencies. The ciliary fragment marker, ARL13B, showed about 1.5-fold increase in conditioned media pellet under hyperosmotic shock (Fig. 8B), while another ciliary protein, IFT88, showed decrease in conditioned media pellet (Fig. 8C). These results demonstrate that a subset of ciliary proteins were released into culture medium by hyperosmotic shock.

**Figure 8.**
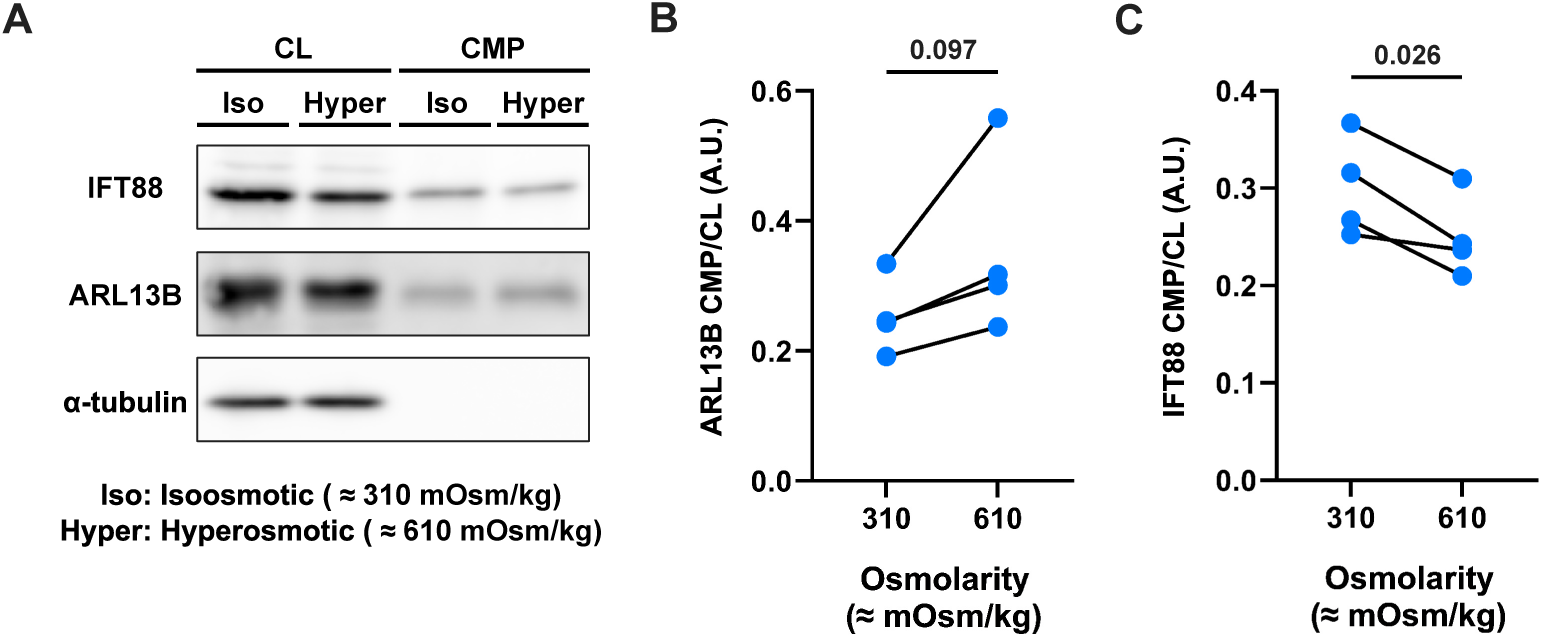
Hyperosmotic shock increases the amount of cilia fragments in the culture medium. **(A)** Western blot analyses for the amount of cilia-related proteins (ARL13B and IFT88) in cell lysate (CL) and conditioned media pellets (CMP) from mIMCD-3 cells cultured at isoosmolarity (≈310 mOsm/kg) or hyperosmolarity (≈610 mOsm/kg) for 3 h. (B and C) Quantification of the ratio of ARL13B (B) or IFT88 (C) amounts in CMP over those in CL as shown in A.

## DISCUSSION

This work demonstrated that acute hyperosmotic shock reversibly shortened and disassembled primary cilia and delocalized PCMs without structural disassembly of the centrioles in the renal collecting ducts epithelial cells. These morphological changes occurred in a manner dependent on excess formation of microtubule and F-actin. Additionally, these changes accompanied the release of a part of ciliary proteins into the culture medium.

We found that primary cilia got shortened/lost with hyperosmotic shock and that the reduction of primary cilia length was NaCl concentration-dependent (Fig. 1). With milder hyperosmotic shock (≈410 mOsm/kg), the primary cilia length was only slightly reduced, and with stronger hyperosmotic shock (over ≈610 mOsm/kg), the primary cilia got markedly shortened and disassembled. This phenomenon seems to be evolutionally conserved as the tonicity-induced Chlamydomonas deflagellation has previously been reported (Solter and Gibor, 1978). An argument could be raised whether the hyperosmotic shock-induced primary cilia shortening/loss are artifacts. The possibility of artifacts can be excluded, though. First, the osmolarity we tried is physiological for the epithelial cells of renal collecting ducts; they are subjected to a very wide range of osmolarity, from 50 to 1200 mOsm/kg (Koeppen and Stanton, 2012). Renal epithelial cells are reported to adapt to these hyperosmotic stresses (Dmitrieva and Burg, 2005). The mIMCD-3 cells we used have been generated by passing through hyperosmotic conditions up to 900 mOsm/kg (Rauchman et al., 1993). In contrast to a previous study where mIMCD-3 cells were exposed to hyperosmotic pressure for a few days (Pihakaski-Maunsbach et al., 2010), the duration of hyperosmotic shock we tested on these cells did not induce cell death in the majority (Fig. 5). Second, our data demonstrate that the hyperosmotic shock-induced morphological changes were reversible (Fig. 4). This reversibility fits well with oscillation of urine osmolarity which depends on water availability and/or intake (Koeppen and Stanton, 2012). Lastly, gradual increase in osmotic pressure also caused the shortening/loss of primary cilia (Fig. 4), which also excludes the possibility of artifacts as urine osmolarity should be altered gradually in vivo. A recently published work further supports our in vitro findings and above-mentioned idea, as water deprivation causes shortening of renal primary cilia in vivo (Kong et al., 2023).

We demonstrated that at least two of the PCMs, γ-tubulin and ODF2, were disrupted with the hyperosmotic shock (Fig. 3), while neither these PCM proteins are degraded, nor nine triplet centrioles are disassembled (Fig. 4). These indicate that the disruption of the pericentriolar protein is caused by delocalization. The PCM is supposed to appear as an electron-dense region in TEM imaging (Dallai et al., 2016). Our TEM data showed that the centrioles had less electron-dense region with the hyperosmotic shock than those with isoosmolarity. This supports the PCM delocalization we showed with ICC and Western blot. The delocalization of the two proteins showed different tolerance against hyperosmolarity; γ-Tubulin was delocalized at ≈710 mOsm/kg or above while ODF2 was delocalized at ≈610 or above (Fig. 3). This difference could result from the different roles of the two proteins. γ-Tubulin is a major component of γTuRCs. γTuRCs are responsible for nucleating microtubules around the centriolar walls (Tovey and Conduit, 2018; Schweizer and Lüders, 2021). The dissolution of γ-tubulin possibly indicates the inactivation of PCM function as centrosomal MTOC surrounding centrioles (Magescas et al., 2019). Contrasting to γ-tubulin, ODF2 is an initiator for the formation of subdistal appendages. The delocalization of ODF2 with hyperosmotic shock suggests the improper forms of distal/subdistal appendages (Ishikawa et al., 2005; Tateishi et al., 2013; Huang et al., 2017). Pericentriolar proteins of PCM are reported to be disassembled after the onset of anaphase at different time points (Mittasch et al., 2020). Different tolerance of γ-tubulin and ODF2 against hyperosmotic shock possibly represents the order of deformation of the basal structure of primary cilia with hyperosmotic shock.

We demonstrated that Nocodazole, a microtubule-disrupting agent, prevented the cilia shortening/loss partially and the PCM delocalization with hyperosmotic shock (Fig. 6). In contrast, PTX, a microtubule-stabilizing agent, failed to prevent the hyperosmotic shock-induced shortening/loss of primary cilia (Fig. S2). These data indicate that the hyperosmotic shock-induced changes depend on excess formation of microtubules by hyperosmotic shock. Several works support this idea. Excess formation of cytoplasmic microtubules seems to limit the tubulin supply for the axoneme in the ciliary space and suppressed the primary cilia elongation (Sharma et al., 2011; Nunes et al., 2013). The result with PTX would mimic the excess formation of microtubules caused by hyperosmotic shock, and thus fail to prevent cilia shortening and loss. Excess formation of bundle-like microtubules could explain the mechanism of a part of PCM delocalization. γTuRCs is reported to also function in a non-centrosomal MTOC (Sanchez and Feldman, 2017). Hyperosmotic shock-induced γ-tubulin delocalization we showed possibly implicates the forceful PCM disassembly accompanying the inactivation of centrosomal MTOC. We assume that the delocalization of γ-tubulin possibly results from diffusion of γ-tubulin to nucleate microtubules extensively in cytoplasm. Delocalization of ODF2 by hyperosmotic shock could result from abnormal interaction between ODF2 and excessively formed bundle-like microtubules, as ODF2 is proposed to have ability to interact with microtubule lattice (Kunimoto et al., 2012).

We found that an actin polymerization inhibitor, LatA, suppressed the cilia shortening/loss partially and the PCM delocalization with hyperosmotic shock (Fig. 7). The results indicate that the hyperosmotic shock-induced morphological changes are actin-polymerization-dependent. F-actin formation activates the transcriptional coactivator YAP/TAZ and inhibits ciliogenesis by expressing AURKA and PLK1 (Kim et al., 2015). Inhibiting actin polymerization is reported to increase ciliogenesis and elongate cilia length (Kim et al., 2010; Avasthi and Marshall, 2012). These imply that the cells increase primary cilia disassembly in an actin polymerization-dependent manner with the onset of hyperosmotic shock. We also showed that the hyperosmotic shock-induced γ-tubulin and ODF2 delocalization was prevented with LatA (Fig. 7). We assume that hyperosmotic shock-induced PCM delocalization occurs in an actin-polymerization-dependent manner. The actin cytoskeleton is reported to control the centrosomes’ subcellular localization by remodeling the actin network and increasing its contractility (Pitaval et al., 2017). It suggests that the cells delocalize the PCM in an actin-polymerization-dependent manner with the onset of hyperosmotic shock.

We detected a trace of primary cilia components in the medium by collecting EV-like particles (Fig. 8). With the hyperosmotic shock, the ciliary GTPase, ARL13B, somewhat increased in the medium pellet. The increased ciliary trace in the medium suggests an increase in ciliary tip scission with hyperosmotic shock. The ciliary tip scission and subsequent cilia disassembly require F-actin formation as faulty F-actin formation inhibits ciliary tip scission (Nager et al., 2017; Phua et al., 2017; Wang et al., 2019). These reports are consistent with our data on increased ARL13B in the medium pellet and suppression of hyperosmotic shock-induced cilia disassembly by LatA. Contrary to ARL13B, IFT88, a component of IFT proteins, was decreased in the medium pellet with hyperosmotic shock. We propose that hyperosmotic shock-induced cilia disassembly accompanies a selective retrieval process for ciliary proteins. This suggests that hyperosmotic shock-induced increased ciliary tip scission and subsequent cilia disassembly are different from the growth stimulation-driven cilia scission where both ARL13B and IFT88 are released into culture media (Phua et al., 2017). The hypothetical process left ARL13B for ectosome cargo and transported IFT88 back to the cytoplasm upon hyperosmotic shock. The loss of axoneme prior to ARL13B-positive ciliary membrane retrieval could explain the hypothetical process (Fig. 2). Shortening of axoneme could sequester IFTs from the tip of primary cilia, while remaining ARL13B-positive ciliary membrane could be pinched off.

Possible biological significance of hyperosmotic shock-induced primary cilia shortening/loss could be argued. In human bodies, hypovolemia makes kidneys concentrate urine. High urine osmolarity and decreased kidney function, lower eGFR, are reported to have a positive correlation (Kitiwan et al., 2021). PC2 in renal proximal tubules, a calcium-permeable cation channel localized on the ciliary membrane, is necessary to maintain normal GFR under fluctuating fluid shear stress (Du et al., 2021). Increased fluid shear stress bends primary cilia, increases calcium signaling from PC2, and eventually increases apical endocytosis in renal proximal tubules. The potential loss of primary cilia with high osmolarity could decrease the kidneys’ capability to adjust GFR causing kidneys to form cysts (Raghavan et al., 2014). The morphological changes of primary cilia and PCM shown in this research suggest that these changes are reversible, temporary cellular responses against osmolarity increase. Primary cilia disassembly accompanying PCM delocalization is possibly triggered to protect primary cilia from fluid-driven shear stress (Renoux et al., 2019) by increased osmolarity as primary cilia have a higher surface-to-volume ratio than cell bodies.

In summary, we have demonstrated that the extracellular stress of hyperosmotic shock shortens and disassembles primary cilia and delocalizes the PCM with the centrioles kept intact in a manner dependent on excess formation of microtubule and F-actin. These morphological changes of primary cilia and the PCM are temporal changes and thus reversible when the extracellular osmolarity is restored. Although the deeper mechanism behind these changes and how they affect cellular functions are still unclear, our findings suggest that the primary cilium is a structure physiologically responsive to extracellular milieu changes.

## MATERIALS AND METHODS

### Cell culture

mIMCD-3 (mouse inner medullary collecting ducts-3) cells (ATCC CRL-2123) were cultured in Dulbecco’s modified Eagle’s medium DMEM/Ham’s F-12 (Wako, Osaka, Japan) with 10% fetal bovine serum (FBS) and incubated at 37°C with 5% CO_2_. Their ciliogenesis was induced by replacing the medium without FBS.

### Generation of Arl13b-Venus mIMCD-3 line

To acquire true knock-in cells (Ijaz and Ikegami, 2019), we inserted Venus into the C-terminus of mouse Arl13B using the HITI method (Suzuki et al., 2016). For CRISPR/Cas9, we used mouse Arl13b 20-bp target sequence and 3-bp PAM sequence (underlined); GCTGTGCGACAGAGACCTAA<u>CGG</u>. To construct Cas9- and gRNA-expression plasmid vector, the 20-bp mouse Arl13b target sequence was sub-cloned into CMV-Cas9-2A-GFP plasmid backbone (ATUM, CA, USA). To construct HITI donor plasmid, exon10 of mouse Arl13b gene tagged at C-terminus with HA and Venus was sandwiched by reversed Cas9/gRNA target sequences and cloned into the TOPO vector backbone. Cells co-transfected with Cas9/gRNA expression vector and HITI donor vector were allowed to grow for 2-3 days and later expanded into a 10 cm dish. Afterward, cells were collected and single-cell-cloned using the limiting dilution-culture method at 0.2∼0.3 cell/well into 96-well plates. Cells were allowed to grow for 1-2 weeks. For the selection of mono colonies, microscopic observations were done to monitor single-cell colony formation and confluency. Selected colonies were expanded into the duplicate of multi-well plates, one of which was utilized to make frozen stocks for subsequent use and the other for screening. Western blot analysis was used to screen the positive clones.

### Hyperosmotic shocks

Cells were subjected to serum starvation by reducing FBS concentration to 0% for 24 h after they reach confluency. We added 5M NaCl to fresh serum-free media to increase its osmolarity from its original osmolarity, rated at ≈310 mOsm/kg. This hyperosmotic media ranges from ≈410 to ≈810 mOsm/kg and the existing culture media was replaced by this hyperosmotic media. PI (Wako, Osaka, Japan) was added for cell death examination immediately after this hyperosmotic shock. To add cytoskeleton-targeting reagent, Nocodazole (Wako, Osaka, Japan), PTX (Taxol, Wako, Osaka, Japan), or Latrunculin A (Wako, Osaka, Japan), we used DMSO as their vehicle (0.1%, v/v).

### Immunocytochemistry

Cells were cultured on nitric acid-washed cover glasses (Matsunami, Japan). They were incubated in each culture media for 3 h and fixed with 4% paraformaldehyde (PFA) in phosphate-buffered saline (PBS, pH 7.5) for 30 min at 37°C. Cells were blocked and permeabilized with blocking solution, 5% normal goat serum with 0.1% Triton X-100 in PBS, for 1 h at room temperature. Then, cells were incubated overnight with indicated combinations of these primary antibodies; anti-ARL13B antibody (17711-1-AP, Protein Tech, IL, USA; 1:400), anti-γ-tubulin (T6557, Sigma-Aldrich, STL, USA; 1:400), anti-ODF2 antibody (ab43840, Abcam, USA, 1:400) and anti-α-tubulin (T9026, Sigma-Aldrich; 1:400) diluted in blocking solution. Cells were washed with PBS and incubated for 1 h with Alexa Fluor 568 or 488-conjugated secondary antibodies (Thermo Fisher Scientific, Rockford, IL; 1:500), Alexa Fluor 488 Phalloidin (Thermo Fisher Scientific; 1:500), and DAPI (DOJINDO, Japan; 1:1000). Fluorescence images were acquired with a laser scanning confocal microscope (Fluoview FV1000; Olympus, Japan or STELLARIS5; Leica, Germany) equipped with an oil immersion lens (60×, NA: 1.35 for FV1000, 63×, NA: 1.4 for STELLARIS5).

### Western blot analyses

The cell lysate was made by adding 1× SDS-PAGE sample buffer to the cells. EV-like particles were collected through a series of centrifugation; 1,000 G for 20 min, 10,000 G for 30 min, and 100,000 G (ultracentrifugation) for 3 h and 10 min. 1× SDS-PAGE was added to make the EV lysate. Either of the lysate samples was heated at 95 °C for 5 min and loaded onto an acrylamide gel. Protein-transferred PVDF membrane was blocked with 5% BSA/TBST for 1 h at room temperature. Then, the membrane was incubated overnight at 4°C with any of the following primary antibodies, anti-ARL13B antibody (17711-1-AP, Protein Tech, IL, USA; 1:1000), anti-γ-tubulin antibody (GTU88, T6557, Sigma, USA; 1:400), anti-IFT88 antibody (13967-1-AP, ProteinTech, USA, 1/5000) or anti-α-tubulin antibody (F2168, Sigma, USA, 1/1000) and diluted in 1% BSA/TBST. Followed by TBST washing, blots were incubated with HRP-conjugated secondary antibodies for 1 h at room temperature and the chemical luminescence was performed by ECL prime (GE, UK).

### Time-Lapse Imaging

Time-lapse imaging was performed as described (Nakazato et al., 2023b). Arl13b-Venus mIMCD-3 cells were cultured in either 35 mm glass bottom dishes (Iwaki, Japan) with DMEM/Ham’s F-12 with 10% FBS. After 48 h of culture, they are replaced with colorless DMEM/F12 medium (Life Technologies Cooperation, USA) for 24 h. In time-lapse imaging, 3-D images are captured every 5 minutes for a total of 3 h using FV1000 confocal microscope equipped with a stage top incubator (Tokai Hit., Co, Ltd. for both microscopes). Before isoosmotic or hyperosmotic exposure, 5 frames of images are acquired, and then medium for isoosmotic conditions (≈310 mOsm/kg) or hyperosmotic shock (≈610 mOsm/kg) is added between the 5th and 6th frame.

### Transmission Electron Microscopy (TEM)

Fixed cell samples for TEM were prepared by the following procedures. Cells were first fixed with 2% glutaraldehyde and 4% PFA in PBS after the hyperosmotic exposure and stored at 4 °C for 2 h, and further fixed with 2.5% glutaraldehyde in 0.1 M phosphate buffer (PB). Fixed cells were washed with 0.1 M PB for 10 min five times at 4 °C, and post-fixed with OsO_4_ in 0.1 M PB for 30 min at 4 °C. The cells were then washed with 0.1 M PB for 5 min twice at 4 °C, and subjected to sequential dehydration with 50%, 70%, 90%, 99.5%, and anhydrous ethanol for 5 minutes each. The dehydrated cells were embedded into a resin, TAAB EPON 812, and the resin was polymerized with stepwise increase in temperature (37 °C for 24 h; 45 °C for 12 h; 60 °C for 60 h). Ultrathin sections with 80 nm in thickness were cut with Ultracut E (Reichert-Jung, Austria) equipped with a diamond knife (DiATOME, Switzerland) at 1 ∼ 1.5 μm above from the bottom of the dish. The ultrathin sections were subjected to negative staining with 3% uranyl acetate for 12 min and with lead citrate for 4 min. TEM images were acquired by means of JEM-1400 (JEOL, Japan) with 80 kV of acceleration voltage and a CCD camera.

### Data analysis

All the data acquired by FV1000 or STELLARIS5 were analyzed on Fiji (ImageJ and associated plugins such as Bioformats) and three-dimensional renderings were created on Volocity (Quorum Technologies Inc., USA). The cilia-positive rate was calculated as (number of cilia)/((number of cells with the whole nucleus in the field) + (number of cells with part of nuclei out of borders of the field)/2). The ciliary lengths were measured by the Pythagorean theorem by using Z projected image and Z slices on which primary cilia propagate. The primary cilia are approximated into straight lines in this method. The statistical significance of each interested pair is evaluated by one-way analysis on variance (ANOVA) on ranks (aka Kruskal-Wallis test) with Dunn’s multiple comparisons tests on the software called Prism (GraphPad Software Inc., USA). Single comparisons were analyzed with Mann-Whitney U test. Paired comparisons were performed with paired *t*-test. A P-value of 0.05 is set to the threshold for statistical significance for our description. The results of the immunocytochemical analysis are representative of three independent experiments.

## ACKNOWLEDGMENTS

We thank Madoka Hamada for her technical assistance and Dr. Qushay Umar Malinta for his support for draft writing. We have no conflict of interest to disclose concerning this study. This work was supported in part by Grants-in-Aid from JSPS for Kiban-C (JP21K06172) and from Japan Science and Technology Agency, Precursory Research for Embryonic Science and Technology (JPMJPR17H1) to KI, and Hiroshima University SPRING fellowship to HO. A part of this work was conducted using research equipment in the Natural Science Center for Basic Research and Development at Hiroshima University (NBARD-00002) under the support from the MEXT Project to promote public utilization of advanced research infrastructure (Program for Supporting Construction of Core Facilities; JPMXS0441300023).

## AUTHOR CONTRIBUTIONS

HO and KI conceptualized the research; HO, RN, and KI designed experiments; HO and RN performed almost all the experiments; KK performed transmission electron microscopy; FI provided Arl13b-Venus-expressing mIMCD-3 cells and prepared EV-like particles; HO, RN, and KI analyzed the data; HO, RN, and KI wrote the manuscript.

**Movie S1.**

The movie shows how primary cilia got shortened/lost during the 3 h of a hyperosmotic shock (right) compared to isosmotic culture media (control, left). Rendered images of representative cilia are shown in Fig. 1F. The cells were stably expressing ARL13B-Venus (red) in colorless media. Time stamps indicate the time after adding the hyperosmotic or isoosmotic medium.

---

**Supplemental Figure 1.**
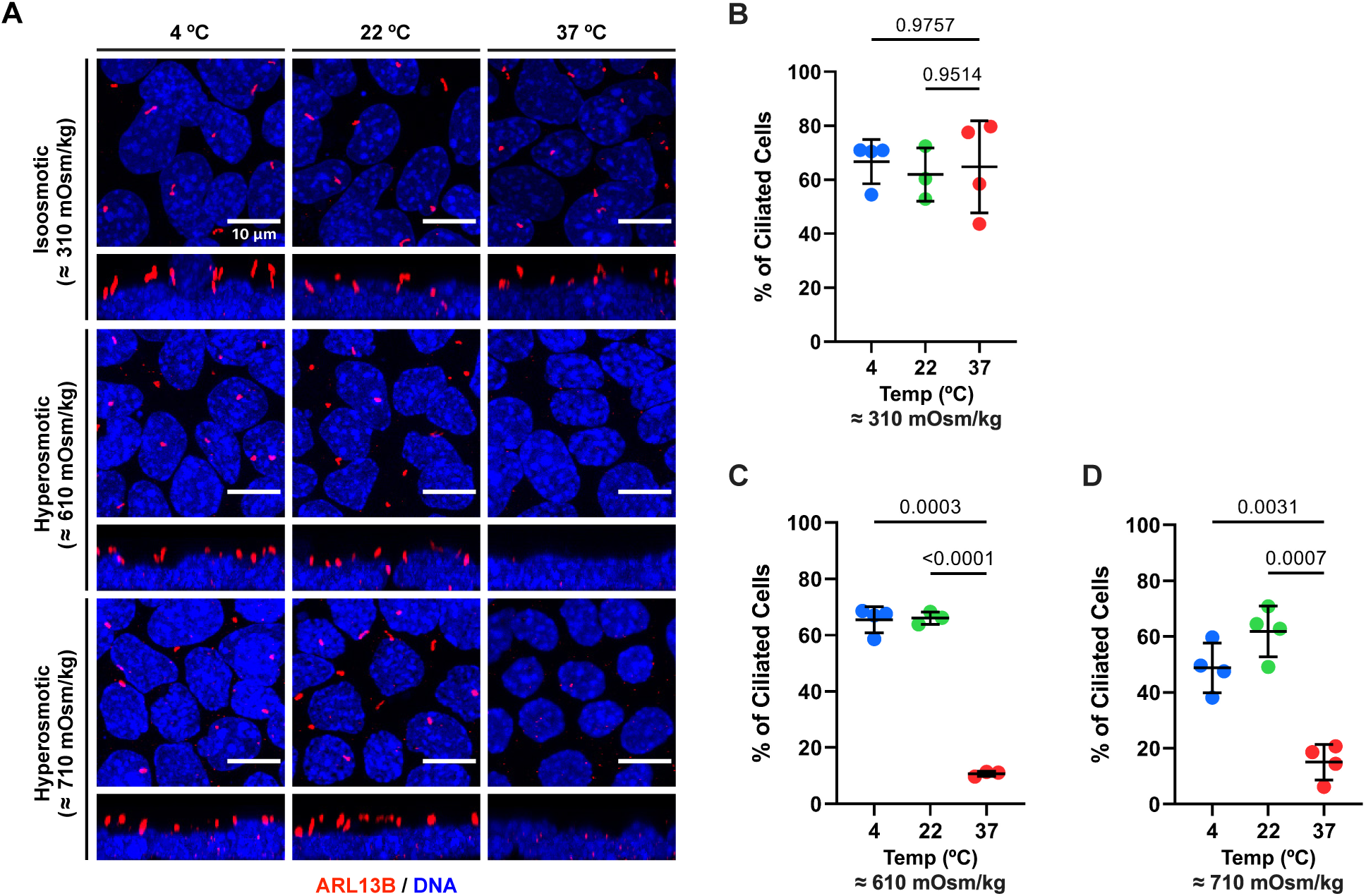
Hyperosmotic shock does not physically disassemble primary cilia. **(A)** Confocal images of mIMCD-3 cells after ICC for primary cilia (red, ARL13B). Cells were cultured under isoosmolarity (≈310 mOsm/kg) or hyperosmolarity (≈610 or ≈710 mOsm/kg) conditions at different temperatures; 4, 22, and 37 °C for 3 h. Scale bars: 10 μm. **(B, C, D)** Quantification of the percentage of ARL13B-positive cilia-possessing cells cultured at different osmolarities, ≈310 mOsm/kg (**B**), ≈610 mOsm/kg (**C**), and ≈710 mOsm/kg (**D**) in **A**. Bars show mean ± SD.

**Supplemental Figure 2.**
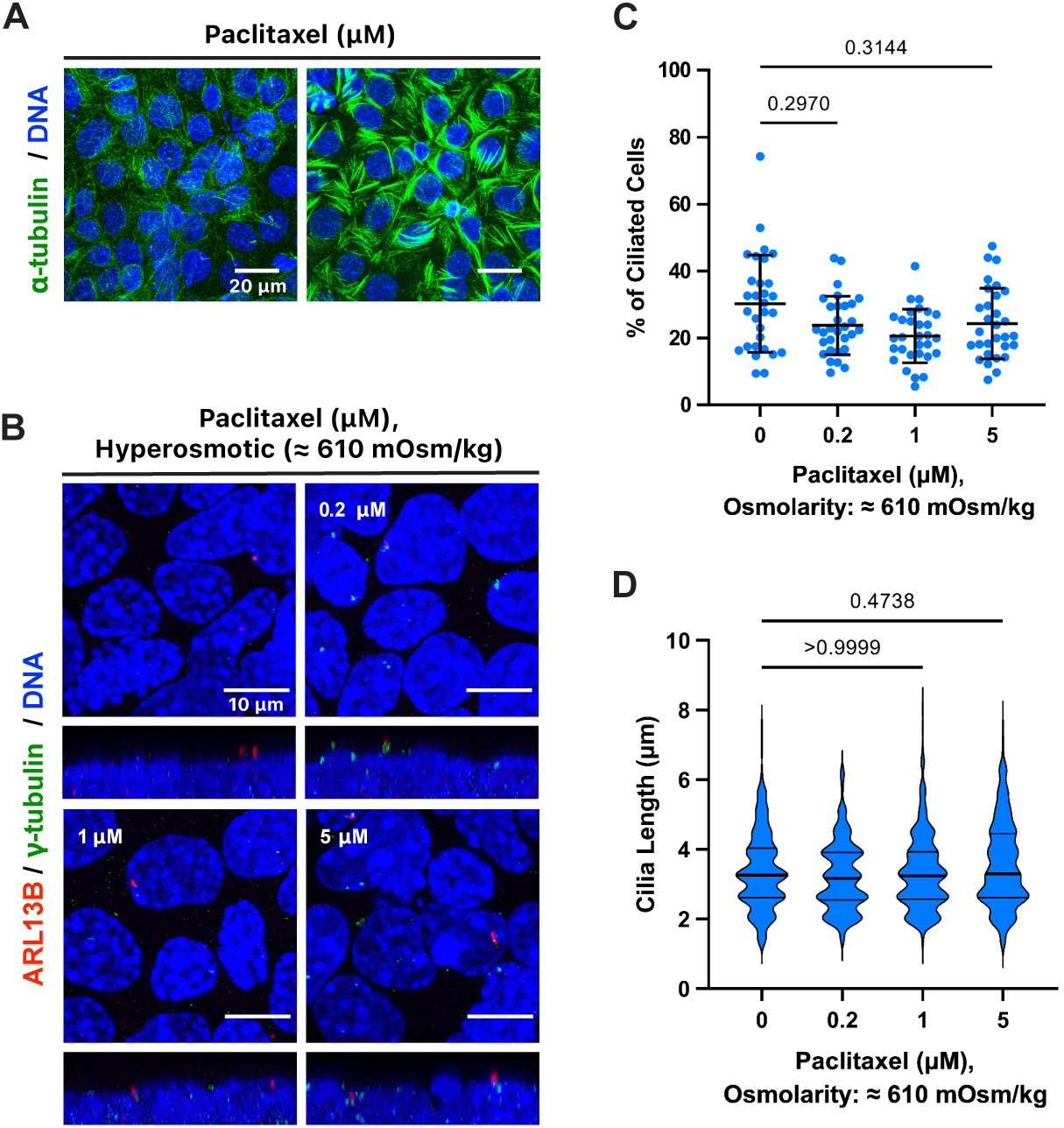
Blockade of microtubule depolymerization did not suppress the shortening and loss of primary cilia upon hyperosmotic shock. **(A)** Confocal images of mIMCD-3 cells after ICC for microtubule (green, α-tubulin). Cells were exposed to the indicated concentrations of paclitaxel (PTX) concentration for 3 h. Scale bars: 20 µm. **(B)** Confocal images of mIMCD-3 cells after ICC for primary cilia (red, ARL13B), the base of primary cilia (green, γ-tubulin), and DNA (blue, DAPI). The indicated concentration of PTX was added to hyperosmotic media at ≈610 mOsm/kg. Scale bars: 10 µm. **(C)** Quantification of the percentage of ARL13B-positive cilia-possessing cells in **B**. The data are from five fields per condition from three independent experiments. Bars show mean ± SD. **(D)** Quantification of ciliary lengths on the cells in **B**. The data are from 10 fields per condition from three independent experiments. Bars show median ± quartile deviation.

